# Prefrontal norepinephrine represents a threat prediction error under uncertainty

**DOI:** 10.1101/2022.10.13.511463

**Authors:** Aakash Basu, Jen-Hau Yang, Abigail Yu, Samira Glaeser-Khan, Jiesi Feng, John H. Krystal, Yulong Li, Alfred P. Kaye

**Author notes:** **Corresponding Author.** Alfred P. Kaye.

## Abstract

Animals must learn to predict varying threats in the environment to survive by enacting defensive behaviors. Dopamine is involved in the prediction of rewards, encoding a reward prediction error in a similar manner to temporal difference learning algorithm. However, the corresponding molecular and computational form of threat prediction errors is not as well-characterized, although norepinephrine and other neuromodulators and neuropeptides participate in fear learning. Here, we utilized fluorescent norepinephrine recordings over the course of fear learning in concert with reinforcement learning modeling to identify its role in the prediction of threat. By varying timing and sensory uncertainty in the formation of threat associations, we were able to define a precise computational role for norepinephrine in this process. Norepinephrine release approximates the strength of fear associations, and its temporal dynamics are compatible with a prediction error signal. Intriguingly, the release of norepinephrine is influenced by time and sensory feedback, serving as an antithesis of the classical reward prediction error role of dopamine. Thus, these results directly demonstrate a combined cognitive and affective role of norepinephrine in the prediction of threat, with implications for neuropsychiatric disorders such as anxiety and PTSD.

## Introduction

Individuals must anticipate and respond to threats in the outside world, selecting from multiple strategies to meet threats along a threat-imminence continuum (Fanselow & Lester, 1988; Moscarello & Penzo, 2022). While a threat far in the future may be met with rumination and intermittent anxiety, an immediate threat will be met with immediate fear and a fight/flight/freeze response across species (Mobbs et al., 2020). However, such behavioral flexibility relies on the accurate computation of threat under varying levels of temporal and situational uncertainty and at various timescales. This objective can be accomplished by the creation of predictive models of the outside world (Peters et al., 2017; Rao & Ballard, 1999).

Threat learning involves the neuromodulation of areas such as amygdala, prefrontal cortex, and hippocampus (Giustino & Maren, 2015; Johansen et al., 2011; LeDoux, 2000; Maren, 2001; Sanders et al., 2003; Takahashi, 2001). Norepinephrine (NE), in particular, has a well-established role in mediating stress (Morilak et al., 2005; Viljoen & Panzer, 2007). For example, NE release enhances sensory perception and increases arousal by modulating forebrain synapses (Berridge & Waterhouse, 2003). NE release is also necessary for classical fear conditioning (Giustino & Maren, 2018; Uematsu et al., 2017). In the amygdala, NE is critical for fear learning and reconsolidation (Bush et al., 2010; Dębiec & Ledoux, 2004; Gu et al., 2020). The prefrontal cortex (PFC) is another important target for NE, with the highest density of NE terminals in the cortex. In the (PFC), NE influences the processing and response to stressors, the expression of fear-related behaviors, and contributes to trace conditioning of fear (Uematsu et al., 2017)(Gilmartin et al., 2014; Giustino & Maren, 2015)(Agster et al., 2013; Arnsten, 2011; Chandler et al., 2014),

Neuromodulators have have computational roles in the prediction of outcomes, although the precise role of specific transmitters is an important area of study. One influential model involves the correspondence of dopamine neuron firing to prediction error in the temporal difference (TD) learning model. In this model, the goal of learning is to predict all future rewards using sensory information within the current state using an estimate of state value. Mismatches between predicted and observed value (i.e. surprise) produce a prediction error (Schultz et al., 1997). Like prediction errors in this model, dopamine neuron firing occurs when there are unexpected, rather than absolute changes in reward value (Schultz, 1998). More recent work has attempted to explain dopamine release dynamics by incorporating the effects of temporal uncertainty to explain dopamine release seen in multistage operant tasks. While dopamine and other neuromodulators are released in response to aversive as well as rewarding cues (Crouse et al., 2020; Hangya et al., 2015; Matsumoto & Hikosaka, 2009; Rajebhosale et al., 2021; Sturgill et al., 2020), the neuromodulatory basis of aversive prediction errors remains poorly understood.

NE has been proposed to have a variety of computational roles in inference, particularly as a representation of uncertainty, salience, and attention (O’Donnell et al., 2012; Yu & Dayan, 2005). NE is known to modulate neural gain across the brain to mediate differential attention to task relevant and irrelevant stimuli, a phenomenon for which multiple models have been proposed (Aston-Jones & Cohen, 2005; Eldar et al., 2013; Mather et al., 2016). There is an activity level-dependent role of NE release in these models, whereby it enhances cognitive function at intermediate levels but produces anxiety and impaired cognition at high levels (Arnsten, 2011). However, technical limitations of prior direct recordings of NE release events limit the ability to understand its precise computational role in aversive learning. The advent of fluorescent sensors of neurotransmitter release has the temporal resolution of NE release measurement, and studies using these sensors found signals related to aversive and reward learning (Breton-Provencher et al., 2022; Feng et al., 2019). In contrast to the precise computational roles assigned to DA which map to reinforcement learning models, a gap exists in the computational understanding of the impact of NE release in relation to aversive prediction. NE has been hypothesized to act as a prediction error signal, particularly during aversive extinction learning (Iordanova et al., 2021). Here, we directly test this hypothesis by combining fiber photometry of neurotransmitter sensors, aversive learning, and computational modeling.

## Results

NE release is necessary for the formation of associative fear learning memories, yet the precise process by which NE mediates the encoding of aversive events in time remains uncertain. By varying the temporal intervals between auditory cues and aversive stimuli (trace conditioning), we examined the role of NE in threat anticipation. NE was measured using in vivo fiber photometry of the GPCR-based fluorescent sensor GRAB-NE2h, which permits selective measurement of NE release dynamics, in the medial PFC (mPFC). These responses were compared during habituation to the sensory stimulus, and then throughout fear learning and extinction. NE dynamics were compared with temporal difference reinforcement learning models of conditioning tasks. Finally, incorporation of temporal event boundaries into reinforcement learning models enabled further investigation of second-to-second NE dynamics during the anticipation of danger. Thus, by combining in vivo NE sensor dynamics with temporally varying danger, were able to interrogate the ability of NE to represent different timescales of threat prediction.

### Prefrontal norepinephrine represents strength of fear association

Dorsal mPFC (dmPFC) facilitates emotional responses to learned fear cues, and this process is facilitated by NE release (Giustino & Maren, 2015, 2018). In order to characterize NE dynamics in dmPFC in relation to fear associations, we combined forward fear conditioning with fiber photometry of the fluorescent NE sensor GRABNE (Figure 1A-B). Freezing increased over the course of tone-shock pairings (Supplemental Figure 1) and was specific to auditory cue periods on Day 3, consistent with extensive work on conditioned associations (Gafford & Ressler, 2011). NE responses to auditory stimulus increased rapidly (Figure 1C) after the first cue-shock pairing and were stable throughout learning (Figure 1D) and continued to be present in the absence of shock in a new context (Figure 1E). NE was not released to the auditory tone in the absence of tone-shock pairings (Figure 1F), suggesting that a sensory salience response cannot explain the cue-induced NE release. These results suggested that the evolution of enhanced NE release might reflect the association strength of a learned cue. Associative learning models, such as that proposed by Rescorla and Wagner, model the associative strength as a gradual accumulation throughout repetition of pairings between a conditioned and unconditioned stimulus (Rescorla & Wagner, 1972). Since we observed such a relationship (Figure 1C-E), we compared the cue-evoked NE release when association strength ought to be zero (Trial 1 Day 1) and when association strength ought to be maximal (Trial 1 Day 3). We found that cue-evoked NE was significantly increased after twenty tone-shock pairings (Fig. 1H), as compared with cue-evoked NE before pairings, which was near zero (Fig. 1G).

**Figure 1:**
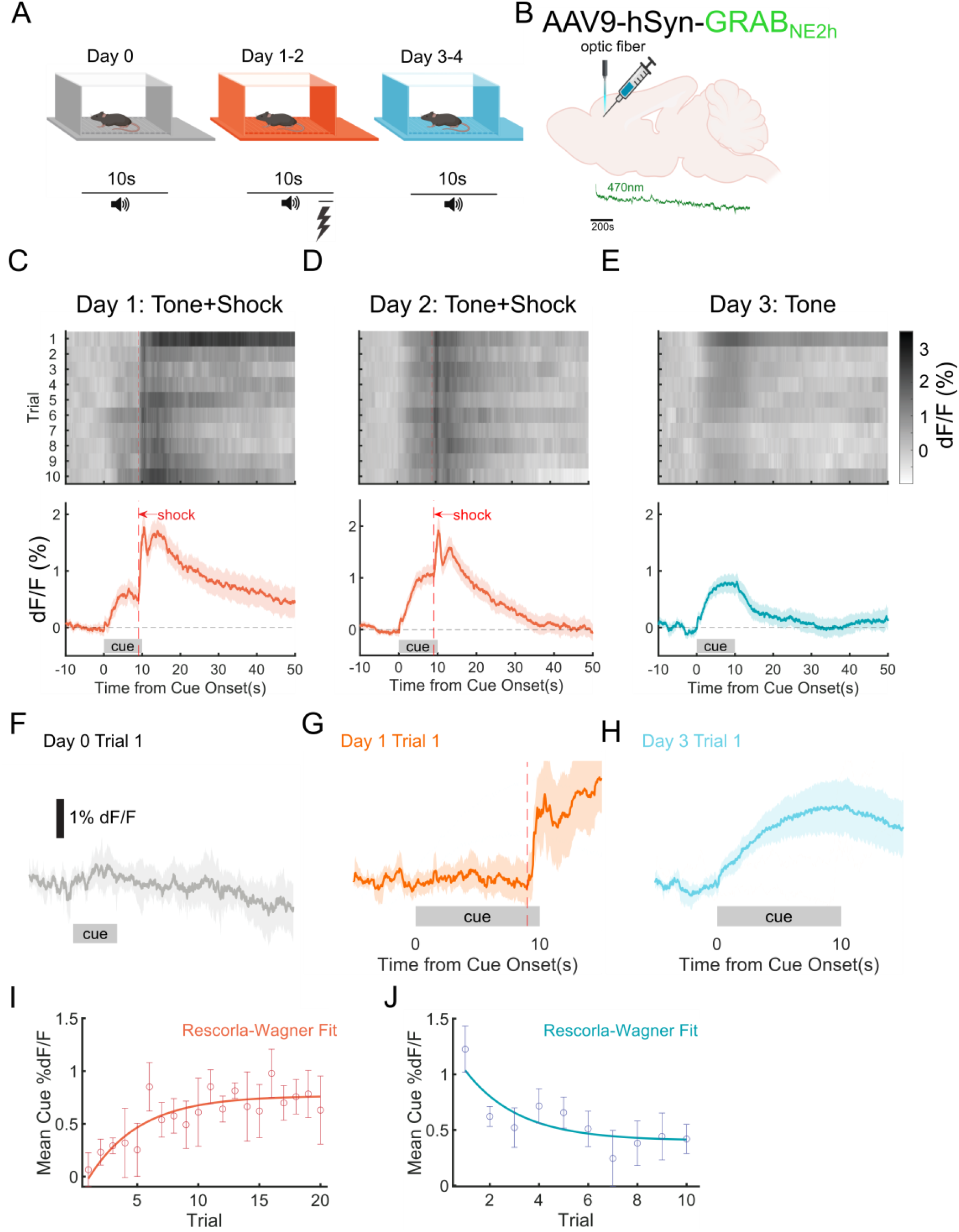
Prefrontal norepinephrine represents the strength of fear association. A: Behavioral schema: Mice (n=12) were pre-exposed to 10 conditioning tones on day 0, then 10 tone/shock pairings (tone and shock co-terminating) per day for two days, then 10 test tones per day for two days. B: Surgical strategy for fiber photometry of GRABNE sensor in the mPFC. C-E: top: trial averages of GRABNE %dF/F for two days of tone/shock pairing (C-D) and one day of tone presentations post training (E). bottom: grand average of %dF/F across trials and animals (10 trials x 12 animals), mean±bootstrapped 95% confidence intervals. F: Cue-evoked norepinephrine release for the first experienced tone without shock pairing. G: Cue-evoked NE release for first experienced tone-shock pairing. H: Cue-evoked NE release for first tone exposure on Day 3, after tone-shock pairing. I: Mean NE release (%dF/F) for the duration of each cue over the 20 tone/shock trials on Days 1-2 (dots and errorbars), and Rescorla-Wagner model fit (curve). J: same as I, but for 10 tone alone trials (day 3).

Associative learning models explicitly predict gradual accumulation of association strength towards an asymptote, which also rapidly declines once the CS-US contingency is abolished. We fit cue-evoked norepinephrine to the Rescorla-Wagner model and found that NE release fit this model both during 20 trials of tone/shock pairing (Fig. 1I, adj. R^2^=0.6869) and 10 extinction trials (Fig. 1J, adj. R^2^=0.6911). Thus, PFC NE release represents the strength of association between a predictive cue and an aversive event.

### Prefrontal norepinephrine shows temporal features of a prediction error signal during trace conditioning

Although cue-induced NE levels closely tracked the strength of fear association, the correspondence between reinforcement learning model components (e.g. prediction error, value, weight) and NE was still ambiguous. In these models, both prediction error and predicted value (Figure 2A) scale with the strength of association asymptotically to a maximum. NE release in response to an aversive event and learned aversive cue (as in Figure 1J-K) are consistent with both interpretations. To dissociate these two possibilities, we introduced a delay between the end of the cue and the footshock (trace fear conditioning, Figure 2A-B). In appetitive learning, striatal dopamine levels ramp upwards between predictive cue end and outcome delivery (Hamid et al., 2015). This observation led to the interpretation that dopamine can represent reward value rather than prediction error. Thus, we addressed whether NE might ramp upwards during the trace using a 15s wait period between cue ending and footshock.

**Figure 2:**
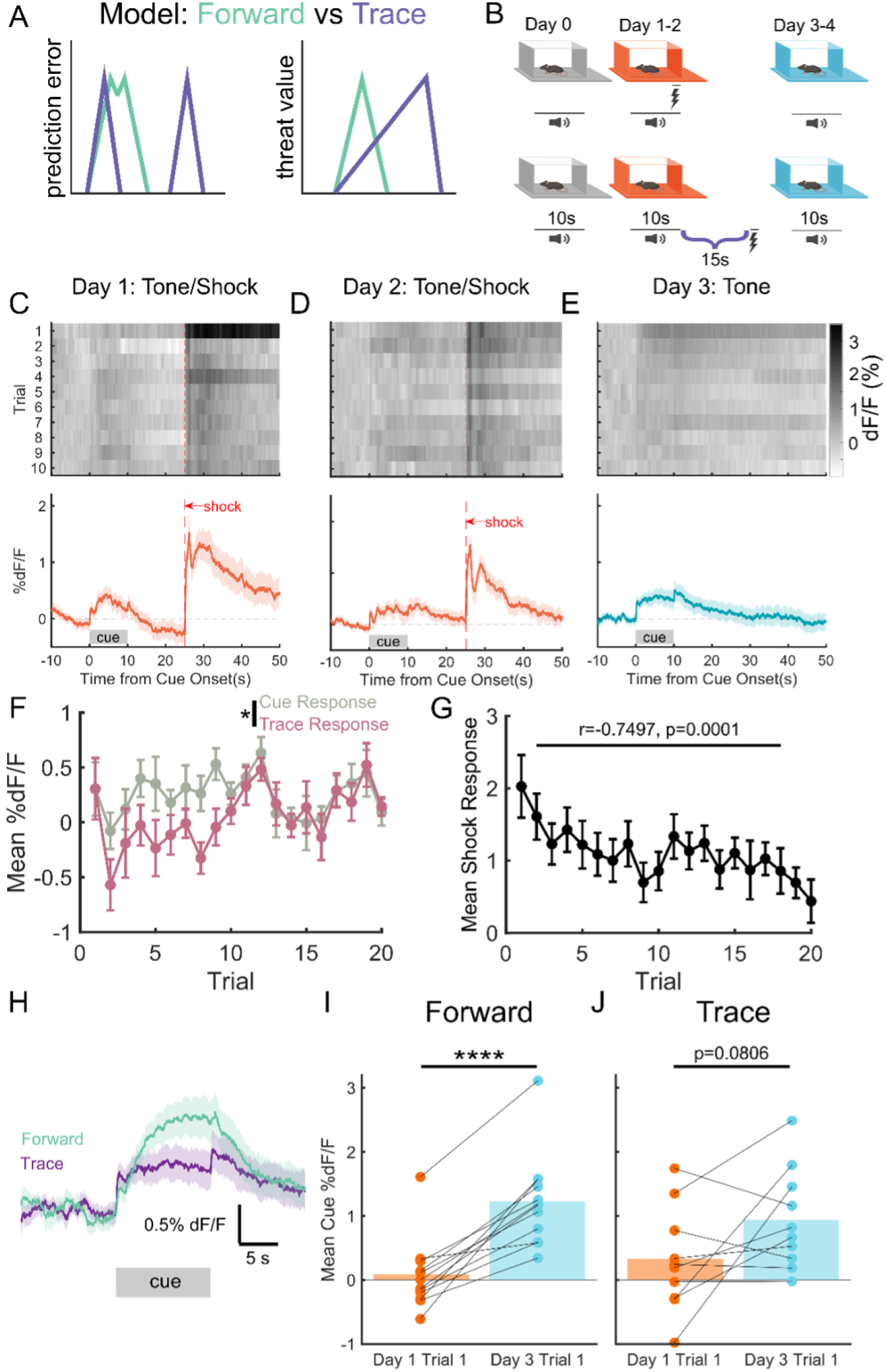
Prefrontal norepinephrine represents a threat prediction error. A: Cartoon of prediction error and value representations. These are difficult to distinguish under forward conditioning due to the proximity of cue and outcome, but with a wait period, prediction error should occur only on the salient events (cue and outcome), while value should ramp between them. B: Behavioral schema. Same as 1A, but with a 15s trace period between cue and shock. C-E: same as 1C-E for 15s trace conditioning (10 trials x 11 animals). Cue-evoked and shock evoked NE are present, but NE release decreases during trace period. F: NE release during trace period is decreased compared to cue period across 20 trials of acquisition. G: Shock-evoked NE release (Shock %dF/F – pre-shock %dF/F) decreases across acquisition trials. H: For cue-only trials (10 trials), cue-evoked NE is lesser for 15s trace conditioning than forward conditioning. I: Forward conditioning increases cue-evoked NE significantly. J: 15s trace conditioning causes significant-trending cue-evoked NE increase.

Consistent with the prior results (Figure 1), we observed cue-evoked NE release during and after training (Figure 2C-E). Critically, after cue-offset, fluorescence returned to near the pre-cue baseline, rather than ramping up to the timepoint of the shock (Fig. 2C-D). Cue onset-evoked NE release persisted in the absence of shocks but habituated over trials (Fig. 2E). A small NE release was visible on the offset of cue in well-trained animals in the presence and absence of shock (Fig. 2D-E), but NE release was decreased during the trace relative to the cue (Fig. 2F). If PFC NE release is indicative of a prediction error, we would also expect that the unconditioned stimulus (footshock) response ought to diminish along tone-shock pairings. Accordingly, we observed a trial-by-trial decrease in shock-evoked NE (Fig. 2G). Thus, the regulation of prefrontal NE levels had features (decrease during waiting period, decreased response to unconditioned stimulus after learning) consistent with a threat prediction error signal rather than a threat evaluation signal.

We observed that cue-evoked NE levels were, on average, lower than that during the forward conditioning procedure (Fig. 2H). As a result, we quantified the ability of NE levels to represent learned aversive cues during trace conditioning as we did for forward conditioning. In forward conditioning, tone-shock pairings significantly increased cue-evoked NE levels between Day 1 Trial 1 and Day 3 Trial 1 (Fig. 2I, t(11)=7.606, p<0.0001). However, we found that 15s trace conditioning produced only a significant-trending increase in cue NE levels between these points of minimal and maximal association (Fig. 2J, t(9)=1.968, p=0.0806). The differential effect of forward and trace conditioning on cue-evoked NE indicated that the ability of NE to represent predictive cues may scale inversely with the temporal distance between cue and aversive event, either due to the difficulty of learning an association between events distant in time or due to a delay-related discounting of the value of the cue.

### Cue-evoked norepinephrine is discounted by trace duration

We observed that cue-evoked NE was smaller in the 15s trace condition as compared to the forward condition (Fig. 2H), suggesting that cue NE is decreased by longer trace duration. In temporal difference models of reinforcement learning, a prediction error update to value ought to be discounted by the time until the outcome (in this case, shock), due to temporal discounting of the value of future outcomes and limited eligibility of a cue in explaining a distant outcome. Thus, if NE release represents a threat prediction error, then the cue-evoked NE release should diminish with increasing trace periods. We tested this hypothesis by varying the length of the trace, using short trace (5s between cue offset and shock) and long trace (30s between cue offset and shock) conditions (Fig. 3A). The trace conditioning cue alone evoked no NE release (Fig. 3B). In the short (5s) trace condition, we observed gradual evolution in cue-evoked NE (Fig. 3C-D) and cue-evoked release of NE in the absence of shocks (Figure 3E. We also observed cue-offset NE release during and after training (Figure 3C-D) as we previously did in medium (15s) trace conditioning (Fig. 2). Moreover, cue-evoked NE was significantly increased by training when comparing the points of minimum and maximum association (Fig. 3F, t(11)=4.385, p=0.0011). However, mice trained with a long trace (30s) lacked any cue-evoked NE across tone-shock pairings and extinction (Fig. 3G-J).

**Figure 3:**
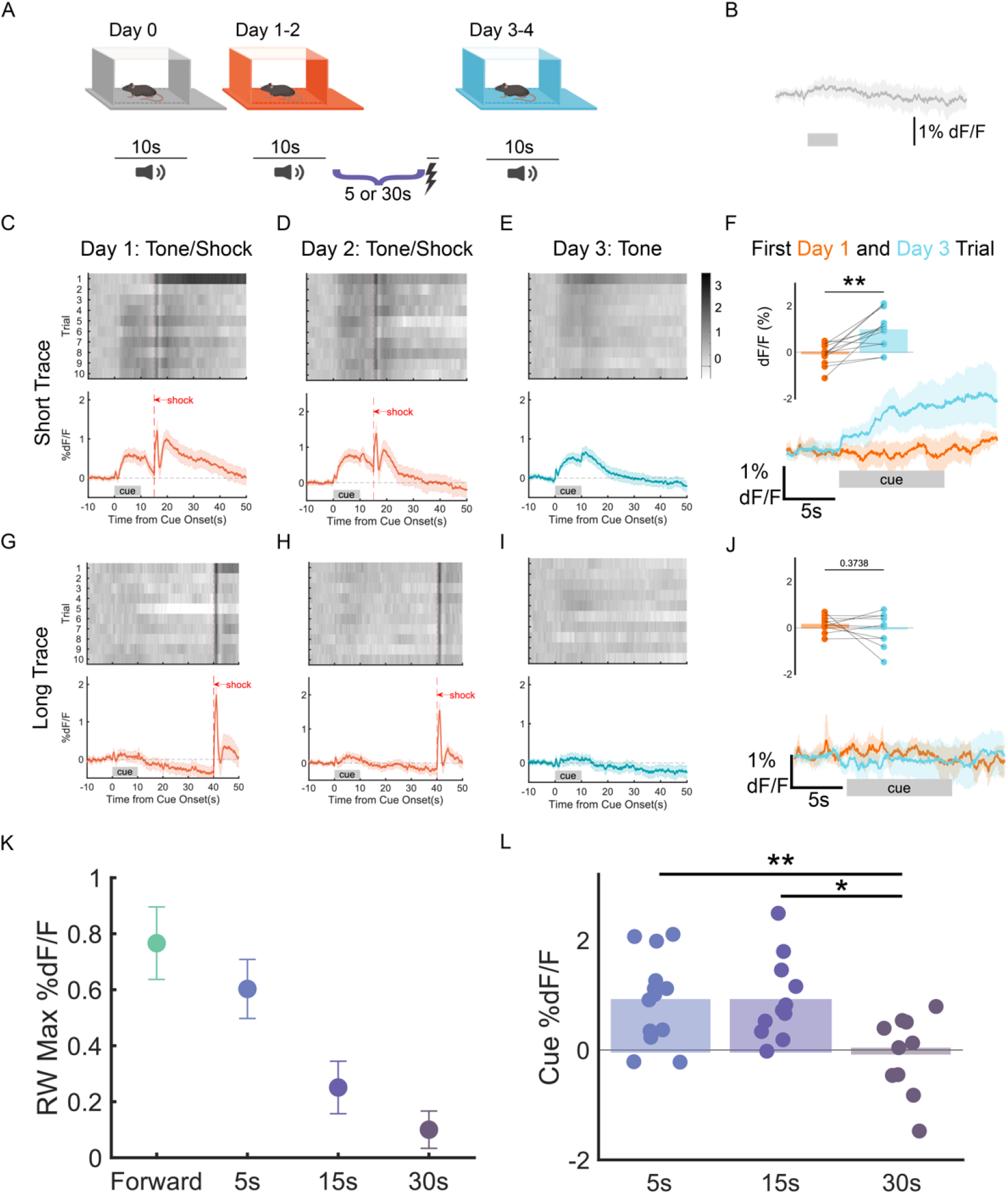
Cue-evoked norepinephrine is discounted by trace duration. A: Behavioral schema: trace duration was varied between 5s and 30s. B: Trace cue without shock pairing evokes no response. C-E: same as 2C-E for 5s trace conditioning (10 trials x 13 animals). F: Cue evoked NE is significantly increased by 5s trace conditioning. G-I: Same as C-E for 30s trace conditioning (10 trials x 10 animals). J: Cue-evoked norepinephrine is not changed by 30s trace conditioning. L: Rescorla-Wagner modeled maximal cue-evoked %dF/F for all conditions. M: Cue-evoked NE is significantly decreased by trace.

Comparing these varying degrees of temporal proximity (i.e. trace length) to threat demonstrated that NE release to a cue shows a decline in maximal association strength with increasing trace length. This decline, as evidenced by the maximum association strength in a Rescorla-Wagner learning model, was gradual for traces ranging from 0-30s (Fig. 3K). Furthermore, when we directly compared the NE release at the maximal association strength (Day 3 Trial 1, immediately prior to any shock omission), we see this same decreasing NE release with trace length (Fig. 3L). NE’s representation of strength of association was diminished by temporal delays, consistent with a prediction error interpretation.

### Prefrontal norepinephrine corresponds to prediction error under temporal difference learning with state uncertainty

The scaling of NE levels with association strength and decay of NE levels during temporal gaps is consistent with prediction error terms of learning models such as Rescorla-Wagner. However, in the noradrenergic representation of trace cue-shock relationship (Fig. 4A), there were some temporal features which could not be immediately explained: the response occurring at the offset of cue (Fig. 4B) and the sustained nature of the cue response (Fig. 4C). In order to apply reinforcement learning models to NE release over time, a representation (complete serial compound representation) of the series of distinct states over time within the trial must be used. Prediction errors can be generated at each of these states by comparing the value at present and future states (Schultz et al., 1997). This approach is used to model temporal dynamics of dopamine release during multi-stage appetitive learning tasks and has yielded insights into the computational properties of dopamine release in relation to task representation in time (Hamid et al., 2015; Keiflin & Janak, 2015; H. R. Kim et al., 2020). However, this model could not reproduce the sustained NE responses observed during the cue period (Supplementary Figure 3A-B).

**Figure 4:**
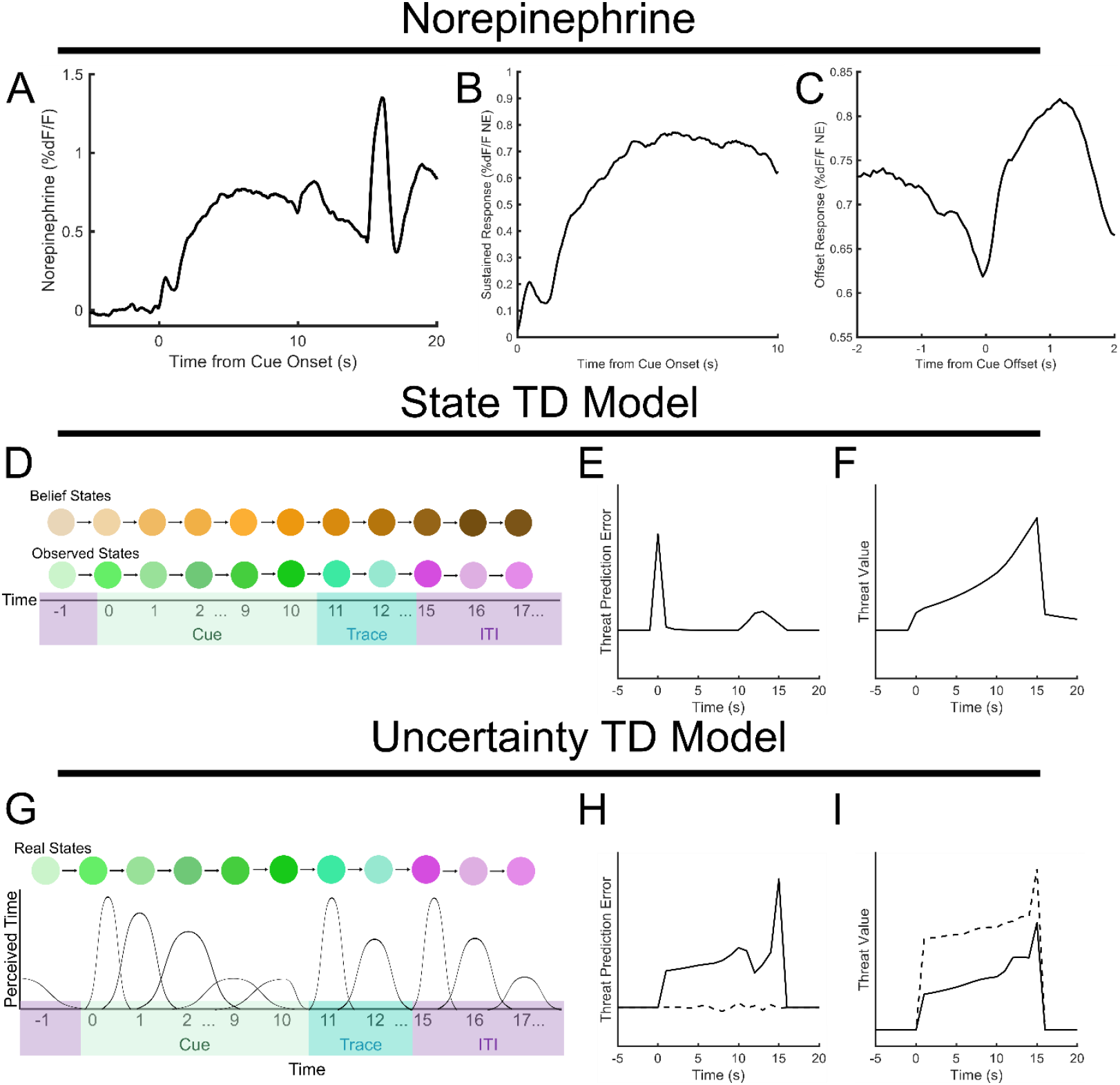
Prefrontal norepinephrine resembles a threat prediction error under uncertainty. A: Mean smoothed norepinephrine release on trials 11-20 (all day 2 trials) of 5s trace conditioning. B: sustained cue response observed during trials in A. C: offset response observed during trials in A. D: Schematic of state temporal difference model: observed states are compared to belief states (from which value is calculated) to obtain a prediction error. E-F: Prediction error (E) and value (F) after 20 trials of training with state model. G: Schematic of state-based TD model with temporal uncertainty. Knowledge of present state is corrupted by uncertainty regarding current time. H-I: Prediction error (H) and value (I) in 5s trace conditioning after 20 trials with uncertainty model. Dotted lines: without correction for uncertainty. Solid lines: with uncertainty correction.

We thus began by modeling our trace conditioning task using a temporal difference learning model in which there could be uncertainty about states (belief states). This model was used to investigate dopamine responses during a trace appetitive conditioning task, and explained variations in dopamine reward prediction errors (Starkweather et al., 2017). In this model, the agent inhabits any of multiple belief states which correspond to the phases of each conditioning trial (cue states, ISI state, ITI state). Each of these belief states is associated with a weight, which is used to compute a predicted threat value (Fig. 4D). This model could reconcile the offset responses we observed during trace conditioning (Fig. 2-3) with a prediction error model, since entering the observable higher threat trace states produces a prediction error (Fig. 4E). In contrast to a value representation in this model, the prediction error showed a lack of ramping in the trace period like our data (Fig. 4E-F). However, this offset response ultimately goes away, which was not observed. Moreover, sustained cue responses could not be generated by this model at the end of training.

One way to generate a sustained prediction error during a sensory cue would be if the subject has an imperfect representation of time. Recently, a reinforcement learning model of dopamine prediction errors due to an imprecise internal clock was used to reconcile the observation of dopamine ramping during appetitive multistage conditioning tasks with the dopamine reward prediction error hypothesis (Mikhael et al., 2022). In this model, value is estimated based on an uncertain perception of time such that states contribute to the value estimate based on their likelihood. When sensory evidence improves the estimation of time (during a sensory cue), the subject has higher confidence regarding which state it is in. Value is therefore continually updated, resulting in a persistent prediction error. In order to apply this model to trace fear conditioning, we assumed that temporal uncertainty would increase with the duration of a time interval, such as cue or trace period (Fig. 4G). Such an increase in temporal uncertainty is predicted by scalar expectancy theory and also observed in the firing properties of time cells (Malapani & Fairhurst, 2002). In such a model, the prediction error term with sensory evidence incorporated, could reproduce both the sustained response to the cue and norepinephrine response to the cue offset (Fig 4H). This was again in contrast to the value representation in such a model, which ramped during the trace period without offset response (Fig. 4I). Intuitively, sustained cue-evoked norepinephrine is produced because each new moment of the cue produces a new resolution of uncertainty and updates value estimates. However, the amount of uncertainty that is resolved decreases throughout the length of the cue, until the offset of the cue, which provides a precise time signal that produces a larger update to value, producing a transient prediction error.

### Time uncertainty model matches learning effects on norepinephrine responses

The NE response to associative cues showed distinct features (Figure 4A) at the onset of the cue, throughout the late part of the cue, and the offset of the cue. These features also evolved differently across learning trials and conditioning paradigms, which we reasoned might define the nature of the reinforcement learning process and thus distinguish between the state model (Figure 4B) and the uncertainty model (Figure 4C). We thus examined key effects of learning conditions (cue onset, sustained cue, and cue offset) on cue-evoked NE responses and compared them to the prediction error terms produced by the models. Longer trace periods produced diminished NE release during the first 5 seconds of the cue (Figure 5A-B). The state model has a learning-related temporal shift which leads to rapidly diminished cue-onset prediction errors in the 15 and 30s conditions at 20 trials of training (Figure 5C), in contrast to the data. This occurs due to a moving prediction error that moves backward in time from shock to cue onset over trials (Supplementary Figure 4A-B), a phenomenon which has been recently observed in dopamine neuron responses to appetitive learning (Amo et al., 2022). Such a prediction error can only “arrive” at the cue onset quickly if the distance between cue onset and shock is short. However, the uncertainty model had consistent responses across the cue-onset period which scaled primarily with the association strength, thus showing a linear decline with increasing trace length (Figure 5D), consistent with the norepinephrine release data.

**Figure 5:**
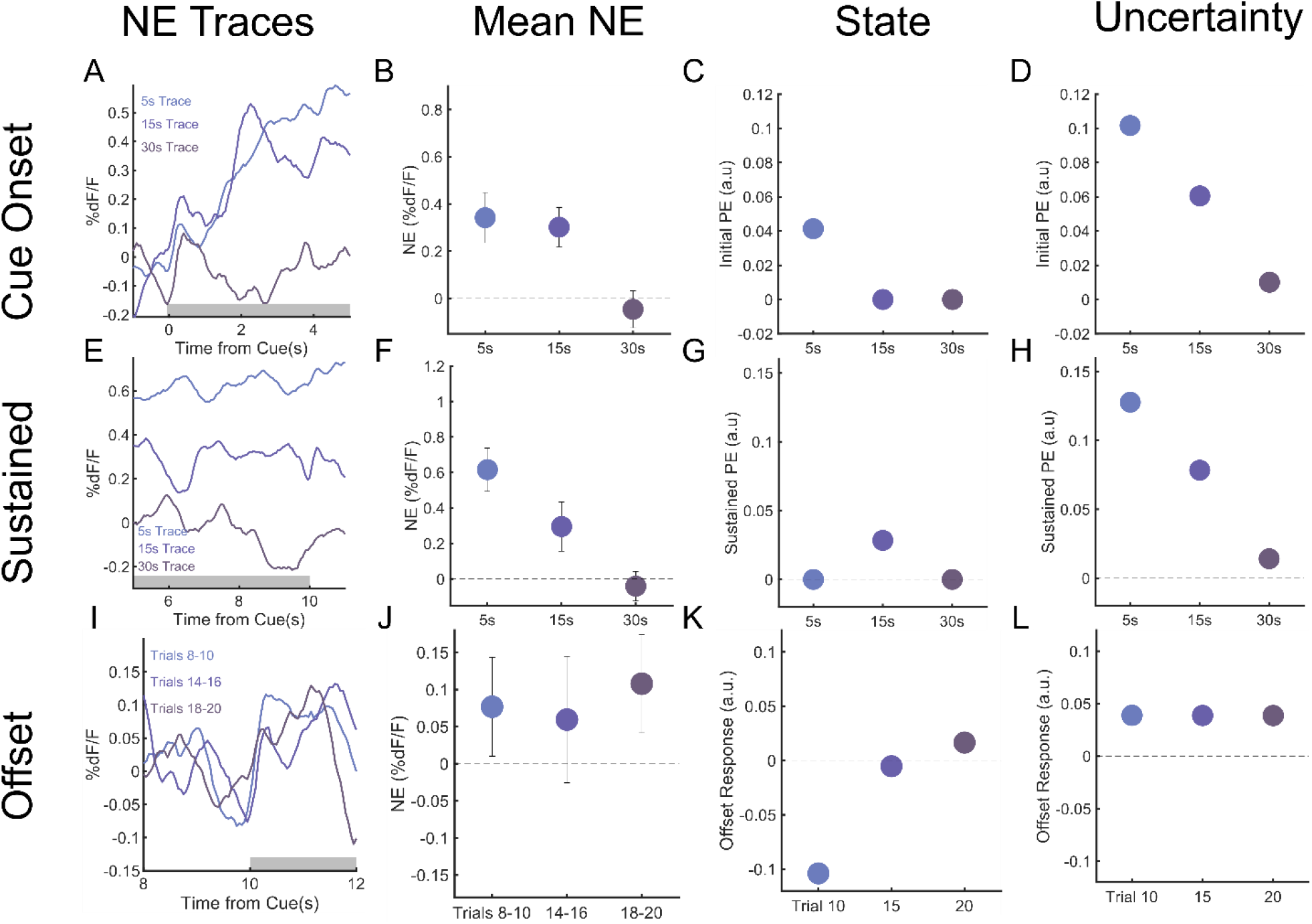
Uncertainty TD model matches learning effects on NE. A: Averaged NE responses to the first five seconds of cue on trials 18-20 of cue shock pairing for 5s, 15s, 30s trace conditioning. B: NE response, averaged along the first five seconds of cue (gray box in A). Mean±SEM. C: Simulated early cue prediction errors for state TD model in 5s, 15s, 30s trace conditioning. D: Same as C but for prediction errors from uncertainty model at 20 training iterations. E-H: Same as A-D, but referring to the last five seconds of the cue (gray box in E). I: Averaged NE responses at the end of cue (10s-12s after cue onset), baselined to end of cue (8-10s after cue onset), for trial blocks 8-10, 14-16, 18-20 of 5s trace conditioning. J: NE responses in I averaged along the cue offset (gray box in I). Mean±SEM. K: Simulated offset responses in the state TD model at trials 10, 15, 20. K: Offsets in the uncertainty TD model at 10, 15, 20 iterations of training. Red lines for all figures indicate 0 on the y-axis.

Cue-evoked NE was sustained throughout the length of the auditory stimulus, which was surprising because in a fully trained subject, no new information is gained after the initial cue onset. This effect could not be explained by convolution with the GRAB-NE2h kernel (Supplementary Figure 5). We observed that the sustained response to the cue, as measured by taking the average NE fluorescence from 5-10s, was decreased by longer trace periods (Fig. 5E-F). This phenomenon is not observed in the state model, where the only sustained response occurs during 15 second trace conditioning due to the temporally shifting prediction error (Supplementary Figure 4). The uncertainty model does capture this feature of the NE release data, since sustained cue responses decreased linearly with increasing trace time (Fig. 5G). The match between data and uncertainty model is due to the discounting of the threat value because of longer trace periods (Figure 5H), since value, and therefore prediction error under uncertainty (Mikhael et al., 2022), decays exponentially with time from aversive outcome. Thus, the uncertainty model, not the state model, captured the trace-induced decrease in sustained response (Fig. 5H).

We observed NE release at the offset of the cue after sufficient training (Fig. 5I-J), which we hypothesized was due to the cue offset indicating more precisely how soon the shock would arrive (by reducing temporal uncertainty). To test this hypothesis, we compared the prediction error terms of both models with NE release over the course of trials. The state model did not produce prediction errors substantial offset responses, except briefly at the end of training (Figure 5K). In contrast, the prediction errors of the uncertainty model produced consistent responses at the offset of the cue throughout learning trials, in keeping with an interpretation of cue offset as a reduction of uncertainty and a transition from the cue to the ITI (Fig. 5L). Observed NE release patterns (Figure 5J) again matched the uncertainty, not the state model, in this respect. Overall, we found that the uncertainty model best explained NE level temporal dynamics over the cue onset (Fig. 5A-D), late cue (Fig. 5E-H), and cue offset periods (Fig. 5I-L), suggesting that noradrenergic representation of threat is modulated by timing uncertainty.

### Cue offset responses are modulated by temporal uncertainty in a long-cue fear conditioning design

If NE offset responses originate from the resolution of uncertainty, increasing temporal uncertainty at the point of cue offset should increase the offset response. Intuitively, if there is very high uncertainty immediately prior to cue offset, the cue offset will produce a large resolution of uncertainty, and thus a large offset response. To test this hypothesis, we designed a variation of 5s trace conditioning where the audio cue is 35s long (Fig. 6A), reasoning that a longer cue should produce greater temporal uncertainty at the end of cue. When simulating prediction errors with the uncertainty TD model, we found that for some parameters, there was a large prediction error at the offset of cue compared to both the 10s cue/5s trace and 10s cue/30s trace variations (Fig. 6B).

**Figure 6.**
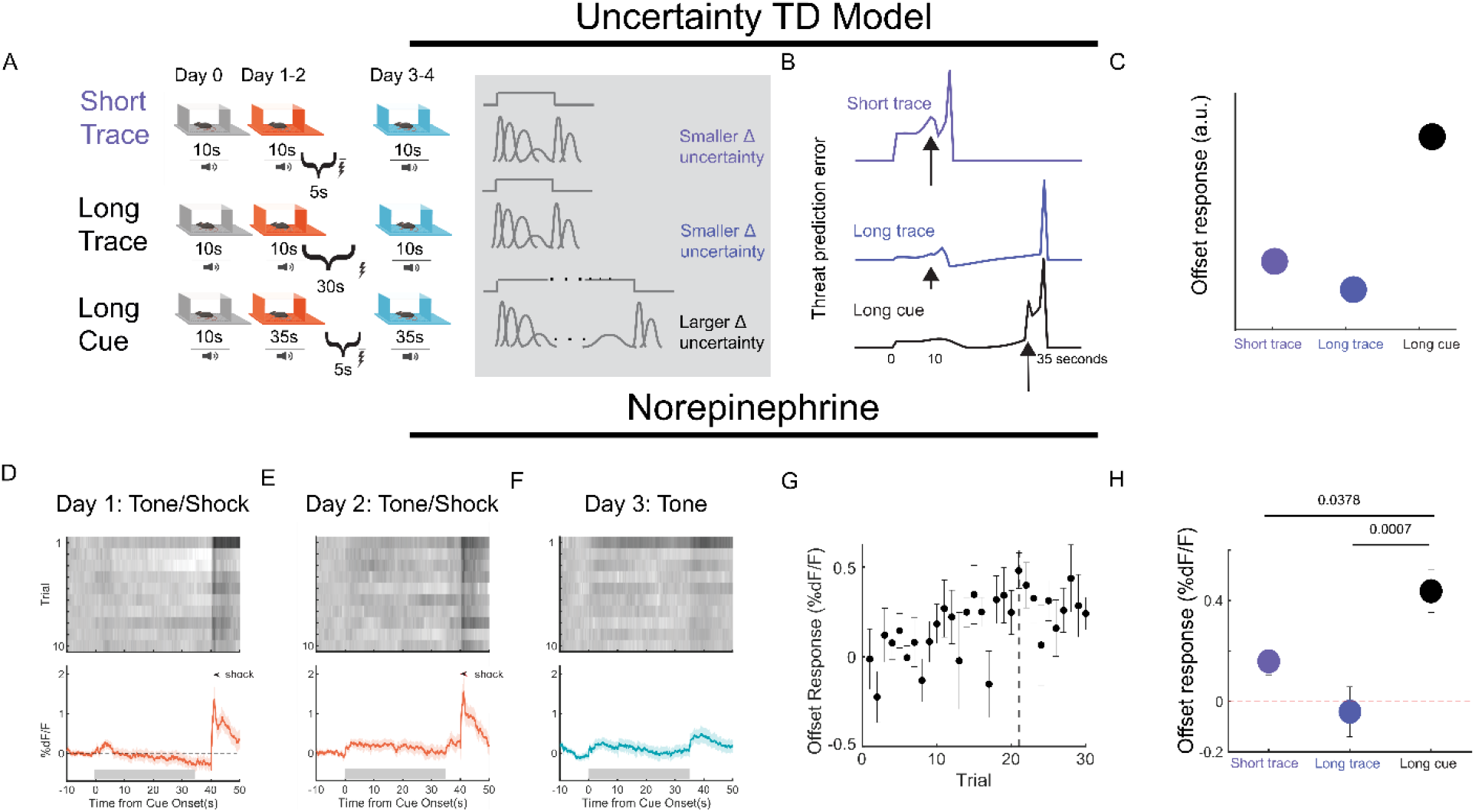
Cue offset responses are modulated by temporal uncertainty. A: Behavioral schema. Animals were trained using 5s trace conditioning, but with a cue 35s long. B: model simulations from 5s trace conditioning, 30s trace conditioning, and 35s cue trace conditioning. C: Offset response in prediction error for short trace, long trace, long cue conditioning simulations with uncertainty TD model. D-F: Same as 3B-D for long cue task. G: Magnitude of offset response (subtracted from end of cue) across training trials and recall trials. H: NE offset responses for 5s, 30s trace and 35s cue conditioning at Day 3 Trial 1: 35s cue conditioning produces significantly higher offsets than either trace task.

We observed that the cue offset response in this version of the task was large when compared to the cue-evoked response (Fig. 6D-F), which existed, but was small compared to 10s cue/5s trace conditioning (Fig. 3B-D). This offset response grew over the course of learning and somewhat extinguished over the course of extinction trials (Fig. 6G). Based on these observations, we sought to compare the offset response observed here to that observed in the 5s trace and 30s trace conditioning experiments. When comparing offset responses at the trial of maximal cue-shock association (Day 3 Trial 1), we found that the offset response from the long cue procedure was significantly larger than both that observed in the 5s trace or 30s trace experiments (Fig. 6H).

### Withholding of sensory feedback eliminates norepinephrine offset responses

Since we had concluded that the NE release at the end of the auditory cue was an update to the animal’s internal clock leading to a prediction error, we sought an experimental manipulation that would diminish this temporal update. In an appetitive visual task, Mikhael and colleagues found that gradually darkening a virtual environment (and thus increasing uncertainty) altered dopamine release in a manner consistent with this model (Mikhael et al., 2022). We hypothesized that gradually silencing the auditory cue would suppress the offset NE release by eliminating the temporal update. Thus, we repeated the short cue/short trace experiment (Fig. 3C-E) but with gradually decreasing volume (Fig. 7A).

**Figure 7.**
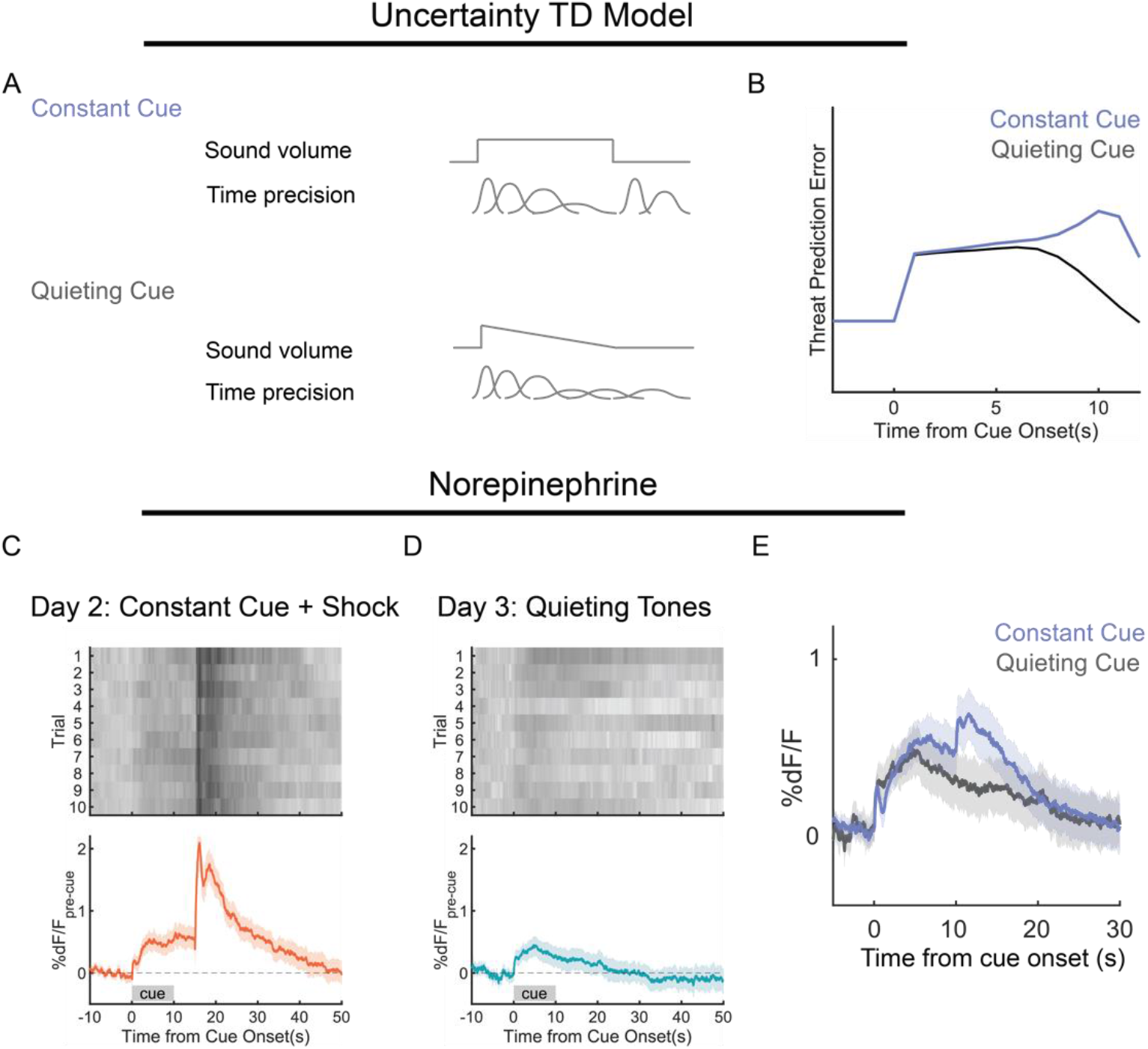
Withholding sensory feedback eliminates offset responses. A: A constant cue will cause a slow decrease in temporal precision, with an update to temporal precision at the end of the cue. A quieting cue will cause faster decay of temporal precision, and cause no such update at offset. B: Model simulation of prediction error with a quickly decaying cue without discrete offset (blue), as well as that for the constant cue as in 4H. C: Trial-by-trial and grand averaged NE response for day 2 of training, with constant tone. D: Same as C, but for day 3 with quieting cues. E: NE responses to constant (Fig. 3E) and quieting cues (as in D) after training.

The temporal uncertainty model predicted that for a fast decrease in sensory feedback over the cue the resulting prediction error should decrease without any offset response (Fig. 7B). In this experiment, animals were trained with the original auditory cue for two days of tone-shock associations, and then the tone was presented with gradually diminishing volume (quieting) on the third day. By day 2 of tone/shock pairing, NE release to the learned cue (Fig. 7C) was unchanged compared with our prior experiment (Fig. 3). However, in the presence of quieting cues, we found that cues exhibited no offset response (Fig. 7D), since there was no abrupt update to the animal’s internal clock at the end of the cue. Like the model, the experimental observation during the quieting cue showed a gradual decrease in NE release without the cue offset increase observed in the constant volume condition (Fig. 7F). Thus, the dynamics of NE release were modulated by manipulating the animal’s sensory feedback about time.

## Discussion

The primary finding of this paper is that NE signaling appears to support the generation of threat prediction errors. Effective responding to threats requires accurate prediction of threats under various timescales and various levels of uncertainty. However, the biological substrates of threat computations have not yet been well elucidated. Here, we sought to determine the computational role of prefrontal norepinephrine release in threat predictions. Using forward fear conditioning, we showed that prefrontal NE release represents the strength of fear association to a learned cue (Figure 1). This is consistent with previous work showing that locus coeruleus firing is synchronized to fear conditioning cues (Martins & Froemke, 2015; Sara & Segal, 1991), and that contextual fear conditioning increases mPFC NE levels (Feenstra et al., 2000). mPFC NE release does not encode stimulus salience or novelty, but only responds to the association of valence to a stimulus. However, this result alone did not fully illuminate the computational role of NE signaling. We used trace fear conditioning to disambiguate a threat prediction error from a threat evaluation, since both reward prediction error and reward value have been hypothesized roles for dopamine release (Mohebi et al., 2019; Schultz et al., 1997). We showed that NE release decays rather than ramps during a trace period between cue and outcome (Figure 2), which contrasts dopamine ramps seen in multi-stage behavioral tasks (Hamid et al., 2015). This observation suggests that NE acts as a prediction error rather than an internal threat evaluation. Had NE served this threat evaluative function, NE levels should have increased progressively between the cue and the shock as animals anticipated the oncoming threat. We also found that cue-evoked NE release during trace conditioning is discounted by trace duration, which is also consistent with a prediction error interpretation (Figure 3).

However, the NE role in threat prediction errors appears to differ from the originally described role of dopamine neuron firing in reward prediction errors (Schultz et al., 1993). We observed sustained increases in NE levels throughout the cue as well as a response on the offset of the cue, in contrast to classical prediction errors which are only observed on cue onset. These observations could be best explained by a reinforcement learning model which incorporates temporal uncertainty (Figure 4). Value estimates of the future are corrupted in a biased manner by temporal uncertainty, and incoming sensory information corrects that bias. In order to compensate for differently computed value estimates in the present and future, there must be a sustained positive prediction error so long as sensory information is present (Mikhael et al., 2022). In our model, we also assumed that the temporal certainty provided by sensory information should decrease the longer the stimulus goes on. We assumed this decrease to be linear, which is supported by the scalar expectancy theory (Malapani & Fairhurst, 2002). This is similar to the idea of microstimuli, a series of stimulus representations which grow weaker and more diffuse over time due to temporal uncertainty (Ludvig et al., 2012). We showed that such a model including temporal imprecision can account for sustained cue prediction errors as well as prediction errors on the offsets of cues (Figure 5). Moreover, similar models can account for the observed persistence of outcome (shock) related prediction errors and absence of negative prediction errors on omission (Gershman et al., 2014; Ludvig et al., 2008). However, we note while sustained dopamine reward prediction errors (e.g. ramps) occur during navigational tasks (Hamid et al., 2015; Howe et al., 2013; H. R. Kim et al., 2020; Mikhael et al., 2022), we observe them here in a classical conditioning task, which may indicate differential modulation of reward and threat prediction errors

This model gave us key predictions about the nature of cue offset prediction errors, an aspect of NE that highlights its role driving adaptive learning in aversive contexts, a role beyond that of NE (especially tonic NE) to produce an anxious state (McCall et al., 2015, 2017; Redmond & Huang, 1979). First, a longer cue should cause a large cue-offset response due to a large resolution of temporal uncertainty as compared to a short cue. We found that NE responses mirror this phenomenon: at the offset of a long cue preceding a trace, the elevation of NE produced by the cue offset was larger than in other conditions (Figure 6). Thus, the offset response is not just a sensory response to the termination of the cue, but is modulated by temporal uncertainty. Conversely, we predicted that in a cue without an offset (a gradually quieting cue), the offset prediction error should disappear. We observed this to be the case for NE release as well (Figure 7). A quieting cue caused NE release at the onset of cue, which decayed through what would have been offset release. Thus, the offset response is not a response to the presence of the trace following termination of the cue, but instead to the uncertainty correction the cue offset provides.

While our observations of norepinephrine release are consistent with a prediction error interpretation, we did not illuminate a causal role for NE release in fear prediction. If NE contributes to generation of a prediction error, it would be necessary for accurate threat predictions. Locus coeruleus firing is both necessary for fear conditioning acquisition (Uematsu et al., 2017) and can drive paired-cue evoked LC firing (Martins & Froemke, 2015), indicating a causal role in fear memory formation. However, these experiments do not disambiguate a prediction error and value interpretation of NE. Future work should determine whether cues paired with evoked NE release would exhibit sustained NE releases or cue-offset responses as observed in cues paired with footshocks. It is also unknown how NE release is modulated by temporal precision. Multiple brain regions such as the prefrontal cortex and hippocampus are known to represent time (J. Kim et al., 2013; Shimbo et al., 2021). It is plausible that these brain regions update temporal precision by projections to the locus coeruleus (Schwarz et al., 2015), but it remains to be determined how temporal precision could be encoded by these projections.

It is also unknown whether NE is a prediction error selective to threat or serves as a teaching signal across valence. Dopamine neuron firing occurs to both positively and negatively valenced stimuli and cues (Kutlu et al., 2022; Matsumoto & Hikosaka, 2009; Vander Weele et al., 2018). Moreover, locus coeruleus firing occurs on a reward predictive cue during a go/no-go task (Bouret & Sara, 2004), In such go/no-go tasks, LC firing occurs upon reinforcement of action, but while expected reward produces heterogeneous firing, unexpected punishment produces strong, synchronous firing (Breton-Provencher et al., 2022). Such observations are consistent with multiple neuromodulators acting as prediction errors across the range of valence. For example, recent work has suggested that dopamine acts as a threat prediction error in addition to its role in reward, and that threat behavior can be best explained by reinforcement learning models which incorporate uncertainty (Akiti et al., 2022). While dopamine has also been proposed to encode salience across valence based on novelty induced dopamine release, we did not observe NE release to non-associated cues, suggesting a bias towards valence (Kutlu et al., 2021), we did not observe significant NE release to a novel, non-associated cue. Acetylcholine has been hypothesized to be a valence-free prediction error which responds to both positive and negative valence (Crouse et al., 2020; Hangya et al., 2015; Rajebhosale et al., 2021; Sturgill et al., 2020). It is likely that an interacting array of neuromodulators produces teaching signals necessary for prediction of incoming outcomes across valence. Future work should focus on the extent to which these neuromodulators are individually necessary in informing predictions, and how they interact to perform predictive computations.

We also observed a tonic component to NE release, specifically on the first footshock or first predictive cue received in a novel context (Fig.1-3). This result could not be predicted by our current TD model, and is less prevalent on trials after the first trial in a novel context. Locus coeruleus neurons exhibit both tonic and phasic firing, with tonic firing varying with arousal state and phasic firing occurring to salient stimuli (Aston-Jones & Bloom, 1981, p.; Berridge & Waterhouse, 2003). It is therefore unsurprising that the first salient cue that animals receive causes both tonic and phasic firing, since these cues must change both arousal state and inform of future events. Future work should determine explicitly whether contextual cues can produce tonic norepinephrine elevations, and to examine behavioral changes produced by elevation of tonic versus phasic NE release.

Our work shows that NE represents a threat prediction error at second-to-second timescales. Norepinephrine is known to mediate stress (Goddard et al., 2010; Viljoen & Panzer, 2007), and modulates cognitive functions such as attention and working memory (Chamberlain & Robbins, 2013). NE’s role in the formation of fear memories has been well established: norepinephrine is released during appetitive and aversive stimuli, modulates excitability of fear-related brain regions, and is critical in fear acquisition (Giustino & Maren, 2018; Sara & Segal, 1991; Uematsu et al., 2017). Previous work has suggested a prediction error role for NE: NE release has been hypothesized to produce the P300 response, which occurs in response to low-probability, salient stimuli, consistent with a surprise or prediction error signal (Nieuwenhuis et al., 2005). Pupil diameter, which is correlated to NE release, has been shown to correlate with prediction errors regarding risk (Preuschoff et al., 2011). However, as of yet a computational role for NE in threat prediction has not been assigned, since this requires measurement of norepinephrine release at fine timescales. Our work seeks to determine this computational role for NE, and assign it a role within a larger framework of threat computation.

Individuals must perform accurate threat computations to choose appropriate behavioral strategies (anxiety, fear, panic) based on the imminence of threat (Mobbs et al., 2020). Analogy can be made between threat computations and reward computations, in which appropriate behavioral strategies must be selected to maximize reward (Sutton & Barto, 2018). While computational models have been richly applied to the neuroscience of reward learning, such models have not received as much attention in threat learning. Associative learning models such as Rescorla-Wagner and temporal difference learning have been applied to threat learning (Cole & McNally, 2007; Levy & Schiller, 2021; Rescorla & Wagner, 1972; Walker et al., 2020, 2022), but biological substrates of components of threat learning models have yet to be found. Our work seeks to begin this process by highlighting one component of a temporal-difference model of threat learning. Moreover, our results indicate that algorithms underlying threat learning are influenced by uncertainty, and that temporal difference models of threat learning should take into account stimulus representations that include temporal uncertainty.

One important aspect of computational models of threat is determining how threat predictions are encoded in the brain. Much work has been done to establish the existence of memory ensembles in the brain, which are activated by fear-associated cues and which are necessary and sufficient for defensive behaviors (Han et al., 2009; Josselyn & Tonegawa, 2020; Liu et al., 2012; Reijmers et al., 2007). Neurons which have increased relative excitability and CREB (a transcription factor downstream of neuromodulators such as norepinephrine) before training are preferentially assigned to fear engrams (Kida et al., 2002; Yiu et al., 2014). How neuromodulators such as NE influence the formation of fear ensembles has not been well studied. If norepinephrine is a prediction error, it should act as a teaching signal to update an internal evaluation of fear. It is likely that norepinephrine prediction errors produce downstream signaling cascades in fear ensemble neurons to encode fear memories. Our work points to a temporally precise role for norepinephrine in the production and regulation of representations of fear memory. Moreover, it suggests a role of NE in the formation of both adaptive and maladaptive fear memories which may occur in disorders such as anxiety disorders and post-traumatic stress disorders.

## Methods

### Animals

A total of 77 animals (45 male, 32 female) were used in this study. Based on histological data, a total of 9 animals were excluded from analysis. Data from two more animals was partially excluded due to recording failures (day 1 data). All experiments used wild-type C57BL6/J background mice purchased from Jackson laboratories at 8 weeks of age. All animals were group housed, and were kept on a 12h/12h light-dark cycle and were provided *ad libitum* chow and water. All experiments were performed during the light cycle (7:00-19:00) All animal procedures were performed in accordance with the protocol approved by the Yale Institutional Animal Care and Use Committee.

### Surgeries

For all stereotaxic surgical procedures, mice were first anaesthetized with isoflurane in oxygen (3-5% during induction, slowly lowered throughout the surgery to 1-2%) while placed in a stereotaxic apparatus (Stoelting). Eyes were lubricated with ophthalmic ointment. Hair was removed with scissors or an electric razor (Phillips), and an incision in the scalp was made to expose the skull. A craniotomy was then made with either a rotary tool (Dremel) or a dental drill above the medial prefrontal cortex. 0.5μl of AAV9-hSyn-NE2h (WZ Biosciences) virus (1 × 10^13^ genome copies per mL) was then loaded into a Hamilton syringe. The tip of the syringe was then placed over the craniotomy and then lowered to the injection site. Injections were targeted to the prefrontal cortex (AP: +1.6-2.1mm AP, ML: ±0.3mm, DV: 1.3mm below dura). Virus was then injected at a rate of 0.1μL/min. After injection, the syringe was left in place for at least 5min, then slowly raised out of the craniotomy. Fiber implantation was done directly after virus injection. Fiber optic implants (0.2mm core, Neurophotometrics) were cut to 3mm length. Implants were held in the stereotaxic apparatus and lowered to the viral injection coordinates. Implants were then secured with quick adhesive cement (C&B Metabond, Parkell).

Mice were then removed from the apparatus and recovered in the home cage. All mice were given lidocaine (5mg/kg, Covetrus) and carprofen (5mg/kg, Zoetis) intraperitoneally after surgery and for two days following surgery. Any behavioral testing took place at least 3 weeks after surgery.

### Behavioral Assays

Mice were handled for 3 minutes per day for 2 days prior to behavioral assays, except for some animals used in experiments in Figure 6 and 7. All behavioral assays were conducted within an aversive conditioning chamber contained in a sound attenuating cabinet (Med Associates). Video was recorded throughout each session using an infrared sensitive camera, and the chamber was illuminated only with a near infrared light. Mice were first placed in an empty cage, and fiber optic cables were connected to the fiber optic implant. The photometry system was calibrated and recordings were started. Mice were habituated to the photometry cable for at least 5 minutes. Mice were then placed into the conditioning chamber, and were allowed to habituate to the chamber for approximately 5 minutes before experimental protocols began.

All experiments took place over the course of 5 days except where indicated. On the first day mice were exposed to 10 presentations of a 10 second cue. For forward conditioning mice the cue was a pure tone of 5.5kHz, for trace conditioning mice the cue was a pure tone of 9kHz. Tone volume was calibrated to 80dBa (as measured from within the chamber) at the start of each experimental day. On the second and third days, the 10 tone presentations were paired with footshocks (1s, 0.5mA). For forward conditioning the tone and shock were co-terminating (shock began 1s before cue offset), for trace conditioning shock began 5s, 15s, or 30s after tone offset. On the fourth and fifth day, 10 tones alone were presented, except for three mice in the 5s conditioning group who did not experience the fifth day. For all experiments, the inter-trial interval (defined as the time between cue onsets) was varied randomly between 60-120s. After tone presentations mice were left in the chamber for approximately 5 minutes before removal. Photometric and video recordings were stopped, and animals were then removed from the chamber and disconnected from the fiber optic cable. Behavioral freezing was then determined from the video file using VFreeze (Med Associates), with parameters adjusted to account for the presence of the patch cord.

Between animals, chamber was cleaned with 70% ethanol. Olfactory cues (lemongrass, eucalyptus, orange essential oil/extract) and visual cues (white plastic insert) were used to distinguish the training context in which shocks were received with the testing context in which shocks were not received.

### Fiber Photometry

Fiber photometry of sensor fluorescence was performed using an FP3002 fiber photometry system (Neurophotometrics). A fiber optic cable (Doric) connected to the FP3002 was attached to fiber optic implants via a ceramic sleeve. Light at the excitation wavelength (470nm) and an isosbestic control wavelength (415nm) was produced from LEDs and propagated through the fiber optic cable to the fiber optic implant. Emission light from the fluorescent sensor was then captured using an sCMOS camera. Excitation wavelengths were used at 10% power (should probably get the real power used), and 470nm and 415nm excitation was interleaved at a total sampling rate of 40hz, producing 20hz recordings for each channel. Only analysis using the 470nm excitation channel was performed, since 415nm is not a true isosbestic point for GRABNE and the dynamic range for the sensor is small (<10% dF/F).

Recordings were saved and analyzed offline. For each cue presentation, the event-related fluorescence change was calculated as a percent change from the pre-cue average. Fluorescence from the 470nm channel was extracted, from 10s before each cue onset to 50s after each cue onset. The percent change for each trial was calculated as a percent change from the average signal from 10s to 0s before cue onset: that is, %dF/F = 100((signal from -10s to +50s from cue onset)-(average of signal from -10s to 0s from cue onset))/ (average of signal from -10s to 0s from cue onset). %dF/F_pre-cue_ was calculated for all trials, and this data was used for all future analyses. Analyses were performed using custom MATLAB scripts, and statistical analysis was performed using MATLAB or GraphPad Prism 9.

### Histology

After all experiments, animals were deeply anaesthetized with isoflurane. Animals were then intracardially perfused with ice-cold 1x PBS, followed by 4% PFA in PBS. Brains were immersion fixed for 1-2 days in 4% PFA, then dehydrated in 30% sucrose. Brains were cut on a cryostat (Leica CM3050S) at 100 micron thickness, and PFC containing slices were mounted onto charged slides. Slides were coverslipped using either Vectashield or ProLong Gold mounting medium. Slides were imaged using the epifluorescence and confocal functions of an Olympus FV3000 laser scanning confocal microscope. Animals were excluded from analysis if viral fluorescence was not detected or a fiber scar could not be localized to the fluorescence.

### Computational Modeling Overview

Reinforcement learning models were adapted from (Mikhael et al., 2022; Starkweather et al., 2017). Temporal difference (TD) models of reinforcement learning attempt to estimate value by incorporating the present value and expected value of future states, discounted by a discounting factor γ. States are assigned weights, and the weight w(b) of the current state b(t) is the value estimate V_h_(t):

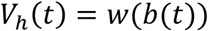

The true value at each step includes any present event r(t) and future expectations of value:

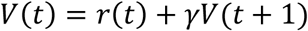

At each step, the difference between estimated and real value is defined as prediction error *δ*(*t*):

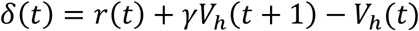

The weight of the current state is then adjusted by the prediction error, scaled by a learning rate:

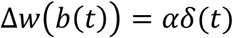

For modeling our norepinephrine data, we used belief-state based TD model in which agents have probabilistic belief states: the agent has a probability of belief that it is within any of multiple states, rather than certainty that it is within one state. All models were constructed in discrete time with timesteps of 1s, and used learning rates and discount factors of 0.9 and 0.9 respectively. In model 1, the value estimate is the weight of all belief states multiplied by the probability of belief of inhabiting each of those belief states;

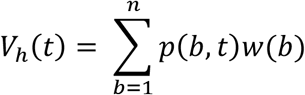

where p(b,t) is the probability of being in state b at time t and w(b) is the weight of state b.

We adapted our uncertainty TD model from (Mikhael et al., 2022). In this model, the agent experiences uncertainty in believed time, and thus temporal state was modeled as a Gaussian probability distribution (uncertainty kernel). Value was computed as an average of state values weighted by the probability distribution p(t). To calculate prediction error, value estimates at the current time (t) and next timestep (t+1) are required. However, in the presence of sensory feedback, sensory evidence causes a narrower uncertainty distribution in the current timestep with standard deviation L(t), but a wide uncertainty distribution in the next timestep with standard deviation M(t). If the value function is convex, wide uncertainty kernels cause value to be overestimated at the next time step, causing biased value estimates in future states and in present states (if prediction error is to converge to 0). Value estimates are thus corrected by a correction factor:

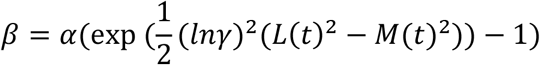

which is applied to the prediction error correction of value. Thus:

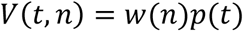

where w is all weights across all time points, and p is the probability distribution of the current time point. Note that p(t) has standard deviation L(t) for t+1, and M(t) for t.

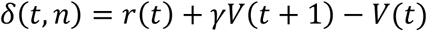

Where δ is prediction error, and V(t+1) and V(t) are calculated based on L(t) and M(t) as above. Value is corrected by:

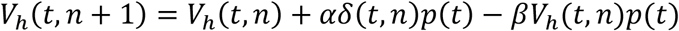

where V_h_(t,n) is the value estimate at time t and trial n. Since value is corrected, δ need not converge to zero and can be positive if L(t) is larger than M(t).

We assumed that for constant cue experiments, the future uncertainty kernels (at time t+1) had a constant, large standard deviation (σ=8s). Present uncertainty kernels (time=t) started with a small standard deviation (σ=0.1s)at the onset of cue, offset of cue, and shock, which were salient events that we assumed would discretely update temporal estimation. With timesteps after these events, the standard deviation increased linearly, with a small increase rate (0.2/s) during sensory evidence (cue) and a large increase rate (2/s) without sensory evidence (ISI). Probability distributions were constrained and then normalized: we assumed that if the animal was in any cue state, it would have no belief that it was outside the cue state series, and similarly for ISI state.

For quantification of model properties, we calculated the initial cue response for both models by averaging the prediction error after 20 trials for states 1 through 5 (first five cue states), and compared to NE cue responses averaged from 0s to 5s after cue onset. We calculated the sustained cue response for both models by averaging the prediction error in states 6-10 (last five cue states) in both models, and compared to NE cue responses averaged from 5s to 10s after cue onset. We calculated offset responses for the state TD model by subtracting a baseline of prediction error signal averaged from state 5 to 7 (late cue), and averaged the resulting signal from state 11-13, since that was where offset responses were observed. For the uncertainty model, we calculated offset responses by subtracting a baseline of the prediction error averaged from states 5 to 7, and averaged the resulting signal between states 10-11, since that was where offset responses were observed. This was compared to NE responses with the average signal from 8-10s (late cue), and then averaged from 10-12s, where offset responses were present.

For model simulations of the long cue experiment, we used a SD increase rate of 0.5 during the cue for all simulations, since that produced an offset response that occurred at the offset of the long cue. For model simulations of the quieting experiment, we used an SD increase rate of 0.5 during the cue, and eliminated the reset to narrow uncertainty kernel at the end of the cue.

## Supplemental Figures

**Supplemental Figure 1:**
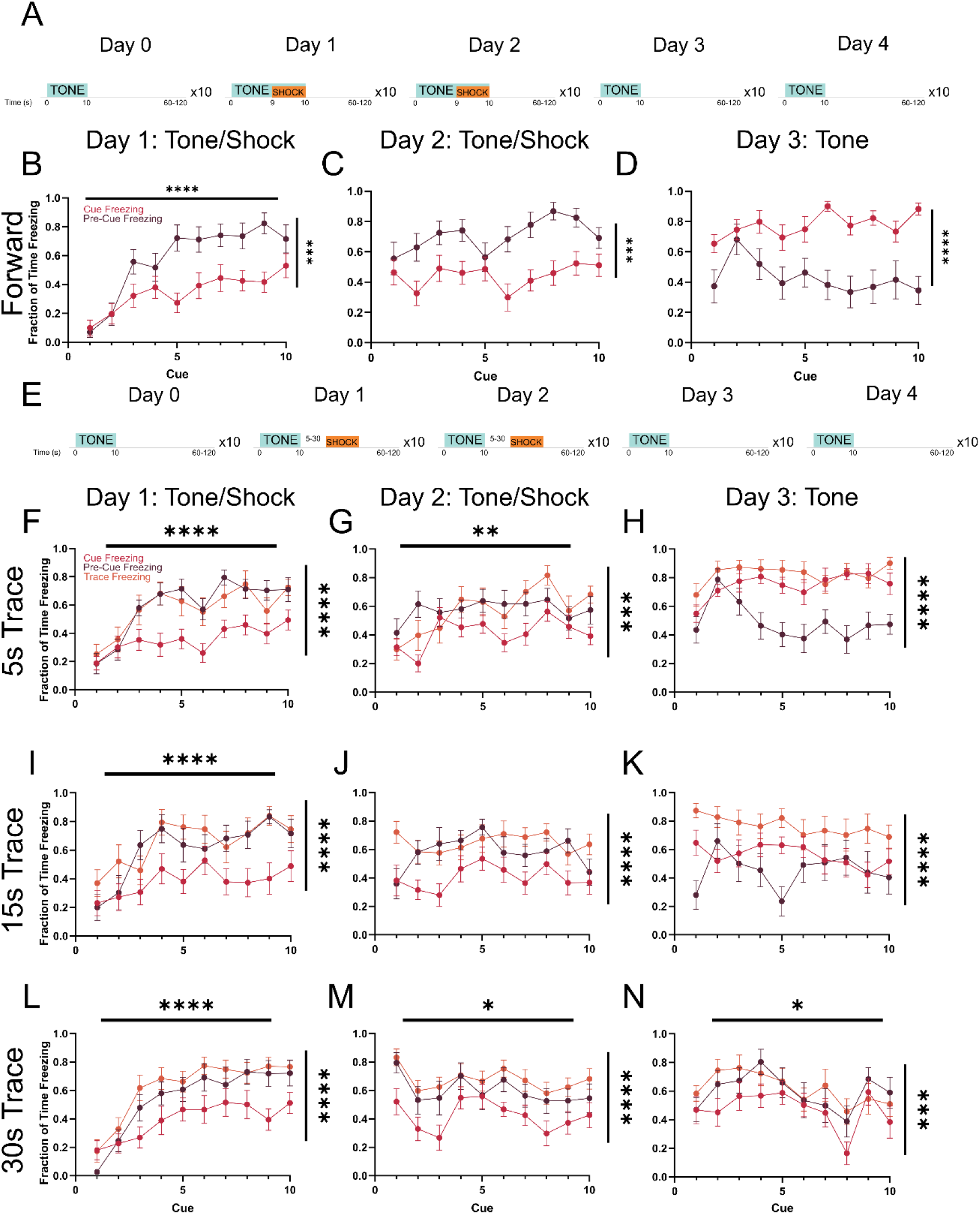
Freezing during forward and trace conditioning. A: Diagram of forward conditioning. B: Freezing in the 10s before each cue (pre-cue) and the 10s of each cue (cue) for initial 10 cue/shock presentations in forward conditioning. C: Same as B for the second day of tone shock pairing. D: Same as C for the 10 tone alone extinction presentations. E: Diagram of trace conditioning. F-H: Same as B-D for 5s trace conditioning, with freezing during trace presented. I-K: Same as F-H for 15s trace conditioning. L-N: Same as F-H for 30s trace conditioning.

**Supplementary Figure 2:**
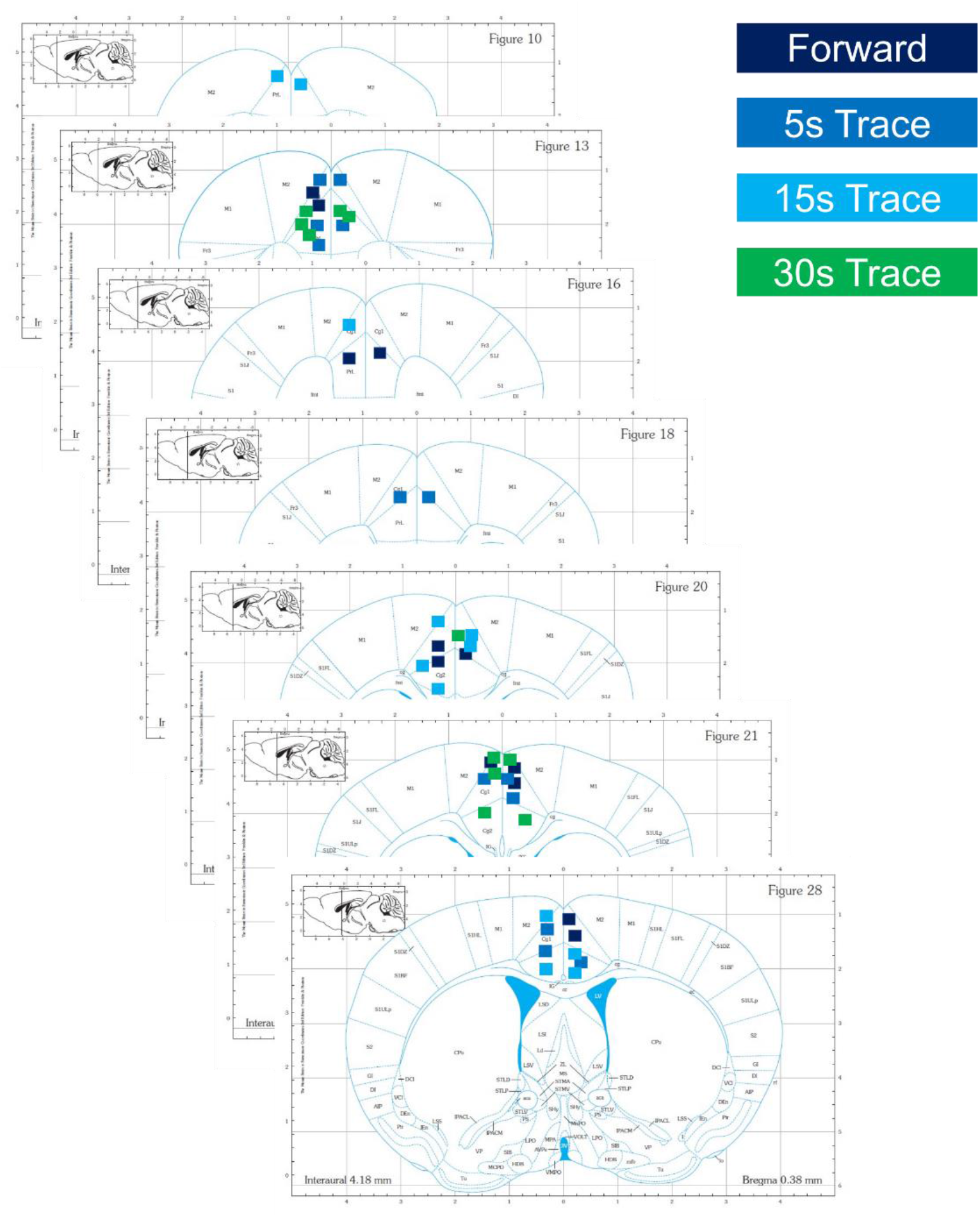
Placement of optic fibers in the medial prefrontal cortex. Rectangles indicate location of fiber, and color indicates experiment of use. Images from (Franklin & Paxinos, 2008).

**Supplemental Figure 3:**
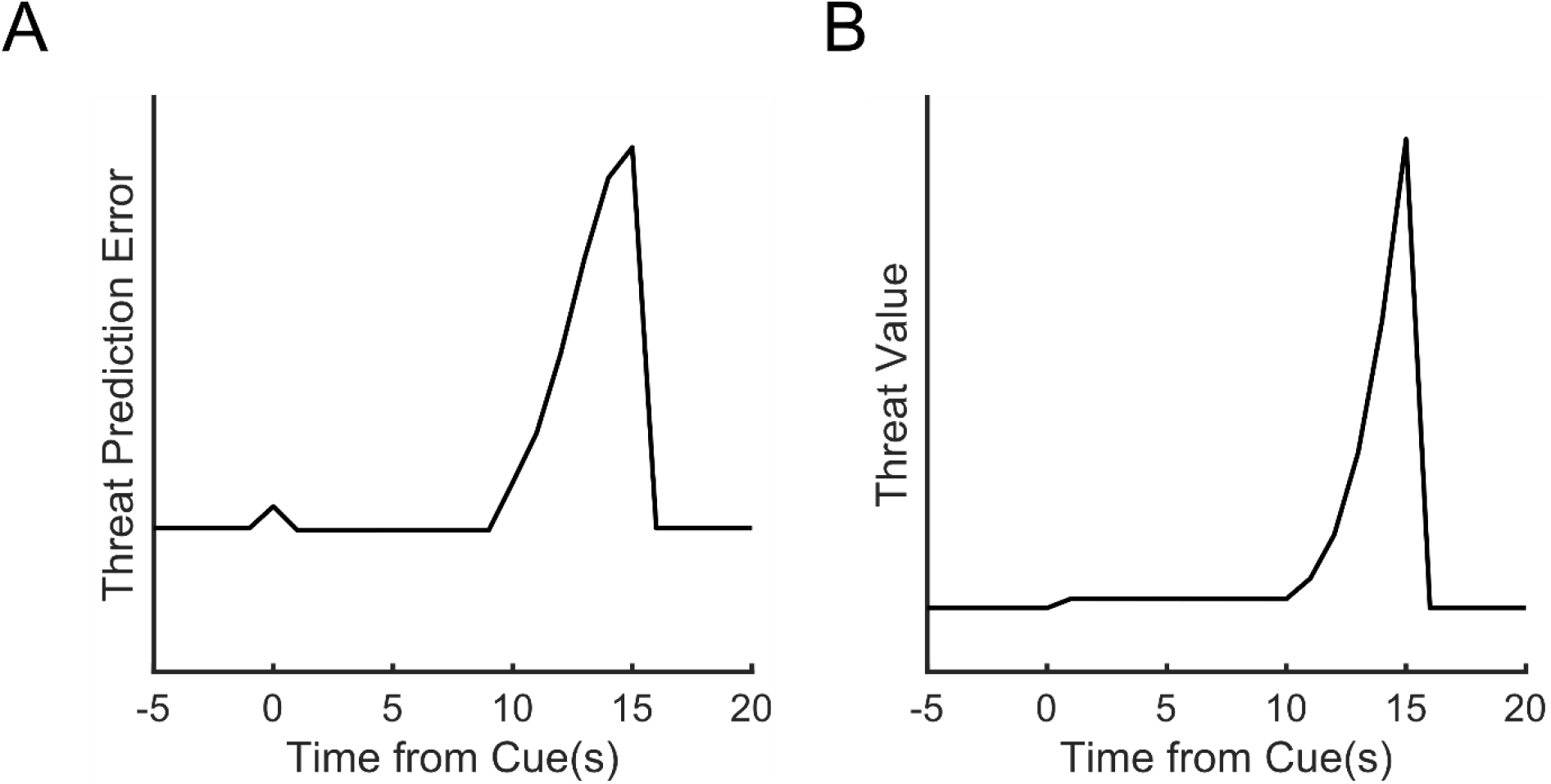
Threat learning modeled using complete serial compound (CSC). A: Threat prediction error after 20 trials of TD learning with CSC. B: Threat value representation after 20 trials of TD learning with CSC.

**Supplemental Figure 4:**
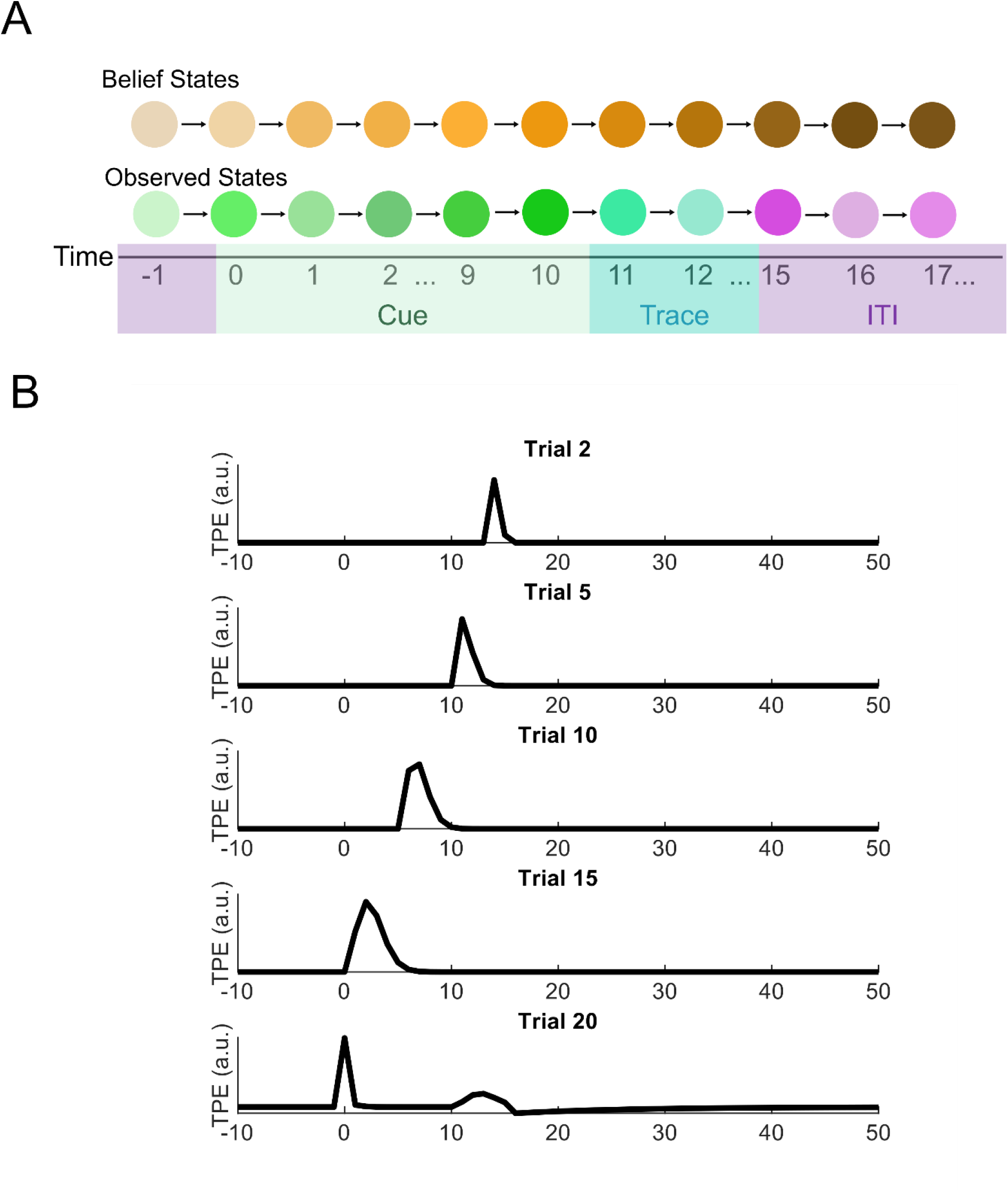
Backwards-moving prediction errors in State TD model. A: Schematic of state TD model as in Fig. 4D. B: Prediction errors across trials move backward along trials.

**Supplemental Figure 5:**
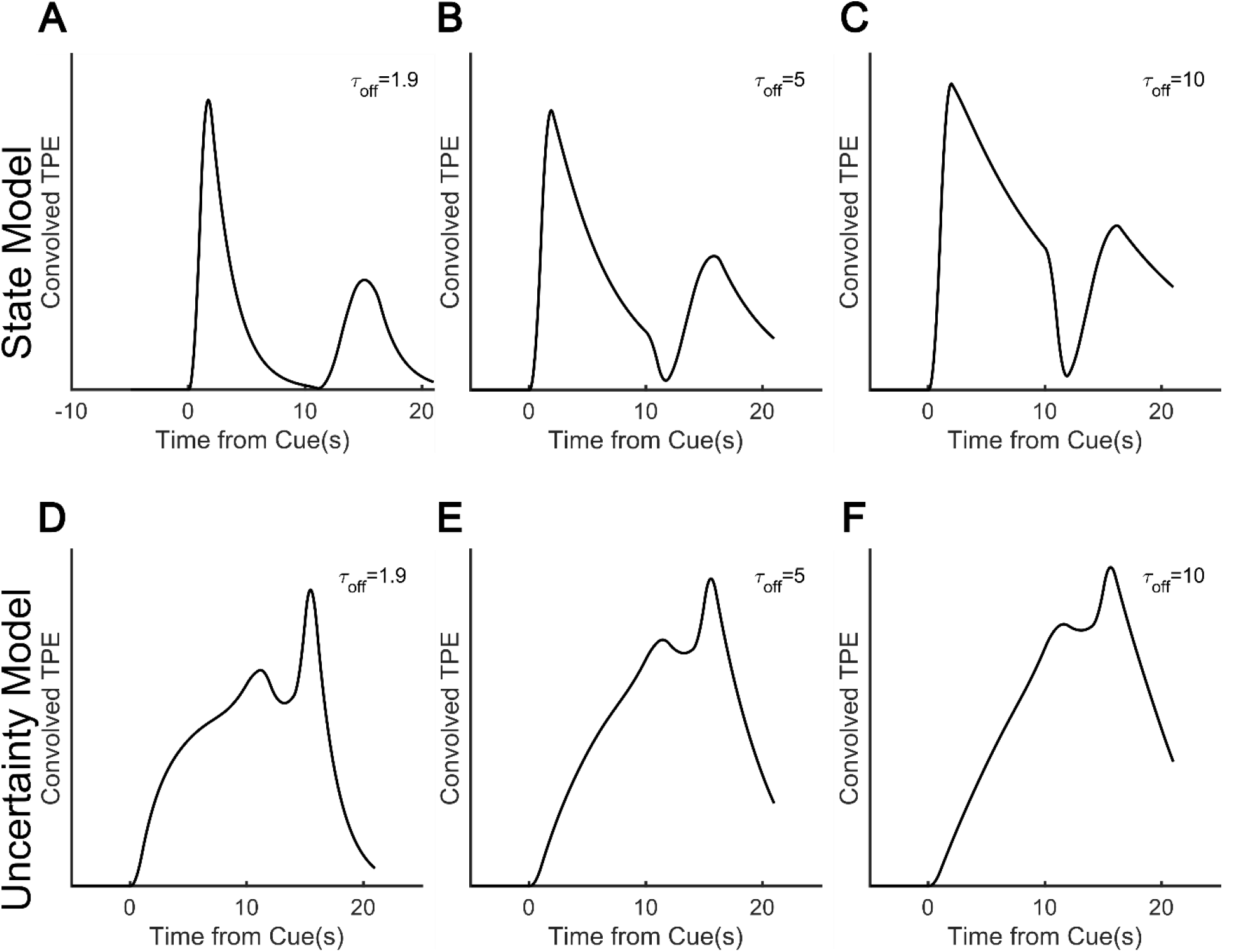
NE sensor kernel width cannot account for sustained norepinephrine responses. A: State TD prediction error convolved with a kernel with off time constant of 1.9s, the off time constant of GRABNE2h (Feng et al., 2019). B-C: Same as A but with time constants of 5s and 10s. D-E: Same as A-C, but using the prediction error from the uncertainty TD model.

